# Consistent differences in eggshell phenotypes select for bluer eggs in an avian host-parasite system

**DOI:** 10.1101/2022.10.27.513857

**Authors:** Juliana Villa, Phillip Wisocki, Jacob E. Dela Cruz, Daniel Hanley

## Abstract

Avian brood parasitism is a well-recognized model for studying coevolution. In this model, hosts adapt the ability to recognize and remove parasitic young to avoid the costs of rearing foreign offspring, while parasites counter-adapt phenotypes to evade detection. A classic example of host-parasite coevolutionary arms race is the great reed warbler and its parasite the common cuckoo (hereafter ‘warbler’ and ‘cuckoo’, respectively). Recent work has found that cuckoo eggs are more likely to be accepted by warblers when those are bluer than their own eggs. Therefore, hosts would select for bluer cuckoo eggs; in turn, this provides directional selection for hosts eggs to remain bluer than cuckoo eggs. Here, we tested whether there were consistent differences in eggshell appearance between cuckoo and warbler eggs using a dataset which provides eggshell coloration of both warblers and cuckoos from the same nest. Despite being a textbook example of mimicry, we found that the cuckoos have significantly browner eggs than the warblers. These findings most likely suggest that this warbler-cuckoo system is experiencing negative frequency-dependent selection, as expected under red queen dynamics. Future research comparing warbler-cuckoo interactions over time would advance our understanding of the coevolution between avian brood parasites and their hosts.

## Introduction

In some ecological interactions fitness advantages for one party result in fitness reductions for another [1,2]. In birds, brood parasitism is one such interaction, which imposes substantial costs on their hosts’ reproductive success [3]. In some systems, such as the common cuckoo (*Cuculus canorus*), the parasitic young even hatches first and kills the host’s entire clutch, resulting in a complete loss of fitness [4]. This explains why many hosts have evolved mechanisms to defend against parasitism by, for example, recognizing a parasite’s eggs or young [5]. When hosts evolve egg recognition, they remove cuckoo eggs they can detect (i.e., relatively poorer matches), which inadvertently selects for mimicry of their cuckoo’s eggshell appearance [6–8]. This, in conjunction with sex-linked eggshell traits, has resulted in the evolution of host-specific races of the cuckoo [4,9]. It is no wonder that this avian host-parasite system is a well-recognized model to studying coevolutionary arms races in the wild [8,10]. These adaptations and counter-adaptations by hosts and parasites can result from a wide range of dynamics other than coevolutionary arms races; however, distinguishing these dynamics within natural populations can be a challenging task.

The *Acrocephalus* warblers are among some of the most well-studied hosts of the common cuckoo [11–15]. Although there are population-specific differences [16], these hosts are adept at egg recognition, and are parasitized by cuckoos whose eggs mimic those of the warblers [16,17]. Generally, hosts are more likely to reject a parasite’s egg if the perceived difference to its own is large [18,19]. More recent work has found that the great reed warbler (*Acrocephalus arundinaceus*) selects for bluer cuckoo eggs by ejecting cuckoo eggs that are browner than their own eggs [15]. Interestingly, this was found even if the absolute perceived color differences between their own egg and both bluer and browner cuckoo eggs were identical. These color-biased rejections would provide directional selection for cuckoo egg colors, and in turn, reciprocal direction on the color of host eggs (i.e., bluer host eggs would have an advantage). Such rejection behavior suggests that there should be consistent differences between host and cuckoo egg colors, such that the warbler eggs should be bluer than the cuckoo eggs. However, relatively few studies have explored such host-parasites dynamics in egg color information for both hosts and parasites from the same nests.

Honza et al. (2014) provided such a dataset that reported individual eggshell reflectance spectra for the eggs of individual great reed warblers and the common cuckoos that parasitized them. This Czech population of warblers rejects natural cuckoo eggs at an intermediate rate (66%); furthermore, this warbler population has historic records of parasitism for at least eight decades [21,22]. These two elements suggest that this warbler-cuckoo community has had sufficient history to co-evolve egg rejection and mimicry. In their study, Honza et al. (2014) found that cuckoos from this population selectively targeted hosts that laid eggs perceptually similar to their own. Studies such as this provide a powerful dataset to test hypotheses that depend on the egg phenotypes of both hosts and parasites, and how co-evolutionary dynamics may be shaping the evolutionary trajectories of both parties [23]. Due to prior work finding color-biased egg rejection in this species [15], we predict that hosts selection for bluer cuckoo eggs would result in reciprocal negative frequency-dependence, such that hosts should have bluer eggs than their parasite. Contrary to expectations under classical coevolutionary arms races, this dynamic would result in parasite and host populations chasing each other’s eggshell appearance, as in a red queen dynamic.

## Methods

### Dataset and model species

We re-analyzed a previously collected dataset of color measurements for 19 clutches of the great reed warbler parasitized by the common cuckoo [20]. These data were collected from May 5 to June 23, 2009 in a fishpond ecosystem between Mutěnice (48º54’ N, 17º02’ E) and Hodonín (48º51’ N, 17º07’ E), Czech Republic. The eggshell coloration was measured as reflectance spectra from 300-700 nm, using a spectrophotometer (USB2000, Ocean Optics). Three different regions along the longitudinal axis of the egg were measured, with three replicates per region, and a total of nine measurements per egg. We used the average reflectance spectrum of each eggshell coloration (see electronic supplemental material from [20]).

### Analysis of phenotypes

We analyzed the eggshell reflectance spectra for the individual host-parasite pairs (n=19) using the package *pavo* in the programming language R [24]; we determined the quantum catches for the ultraviolet, short, medium, and long wavelength photoreceptors under the average avian UV visual system [25]. Based on these quantum catches, we calculated the coordinates of each egg within a tetrahedral color space [26] and estimated the perceived chromatic and achromatic contrasts in units of just noticeable differences (henceforth ‘JNDs’) between the robin egg and each warbler and cuckoo egg. We then transformed the JNDs to directional JNDs multiplying by 1 the JNDs when they were equal or greater than the robin egg contrasts, or by -1 when these JDNs were lesser than those of the robin [27]. Then we constructed a phenotypic space based on the directional JNDs for the avian perceived color and brightness [28,29]. Finally, we calculated the angles and Euclidean distances between each host, parasite, and a common reference point (Fig. 1). If the great reed warblers tend to reject browner cuckoo eggs than their own, we would expect warblers and cuckoos adapting bluer egg colors over time [15]. Thus, we used the eggshell reflectance of the blue American Robin (*Turdus migratorius)* egg as a reference in this model (see electronic supplementary material).

**Figure 1.**
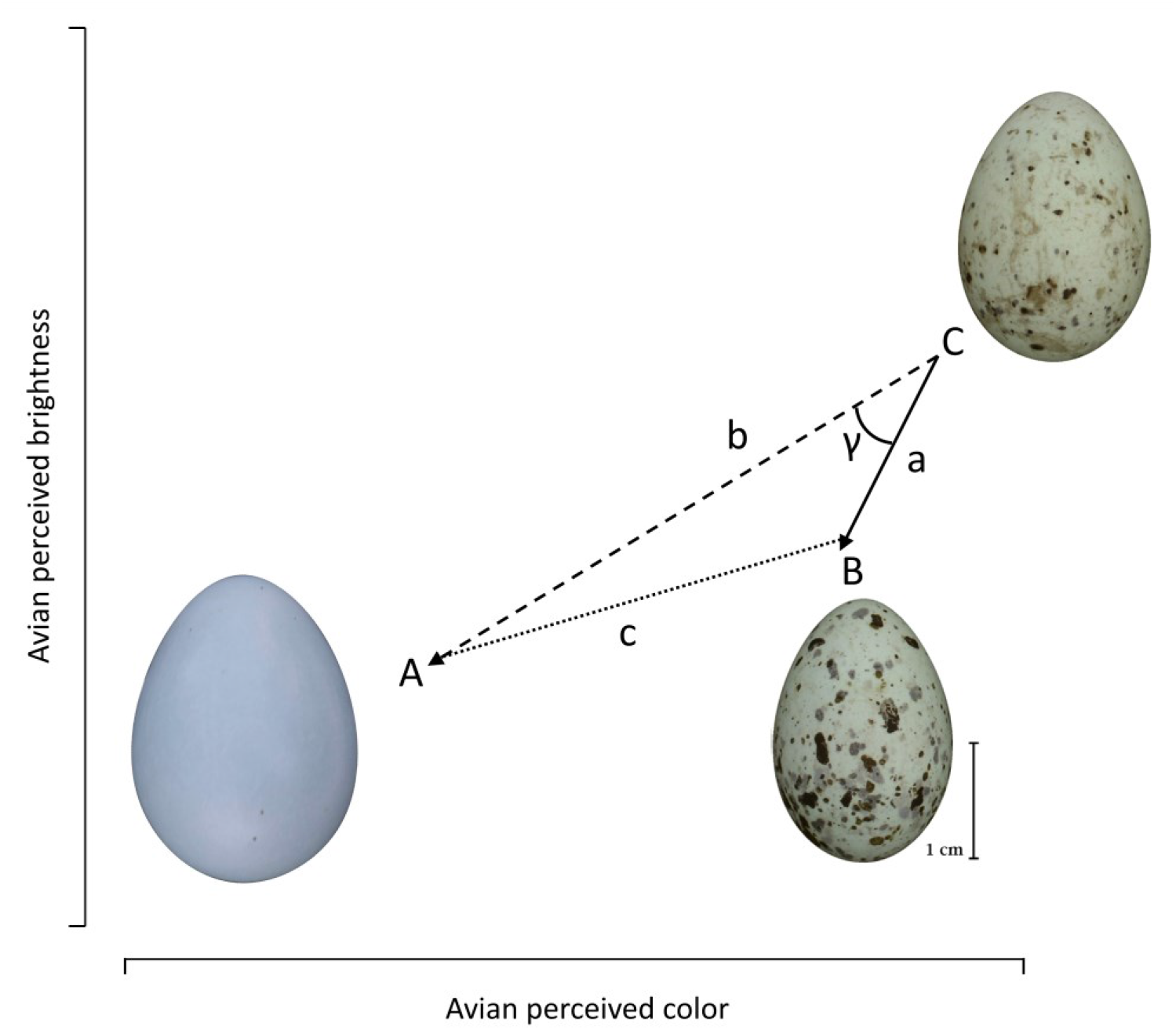
Euclidean distances between the eggs of the American robin (A), the great warbler (B) and the common cuckoo (C) within a phenotypic space where the avian perceived color and avian perceived brightness represent respectively the x- and y-coordinates. To calculate the angle *γ* between the individual host-parasite eggs, we determine the phenotypic distance between cuckoo and warbler (vector *a*), warbler and robin (vector *b*), and robin and cuckoo (vector *c*).

### Statistical analysis

We used circular statistics to analyze and visualize the angle of a vector between pairs of warbler-cuckoo eggs within the phenotypic space, using the package *circular* in R [30]. We then used a Rayleigh’s test to determine whether these angles were random or uniformly distributed. This statistic relies on a parameter *r*: *r* values close to zero represent random directions and *r* values close to one indicate uniform directions. If there is no consistent phenotypic direction between the host and parasite egg phenotypes (i.e., random dispersion), we would expect a uniform distribution of the vector directions, and an *r* value close to 0 (i.e., null hypothesis) [31]. In other words, a parasite’s egg could be bluer or browner, and darker or lighter, than its chosen host. Conversely, if there was a consistent phenotypic direction between the host and parasite egg phenotypes, we would expect a nonuniform (i.e., nonrandom) distribution of vector directions, and an *r* value close to 1 (i.e., alternative hypothesis) [31]. The latter hypothesis would suggest that the parasites are consistently either bluer or browner, or darker or lighter than their hosts within the phenotypic space.

## Results

The trigonometric approach revealed that the avian perceived color and brightness between cuckoos and warblers are consistently different in the Czech system (Fig. 2). Specifically, we found that the warbler eggs are significantly bluer (*t* _(18)_ = -3.896, *p* = 0.001; mean ± s.e.= *warbler*: 5.649 ± 0.138, *cuckoo:* 6.597 ± 0.249) and darker (*t* _(18)_ = -12.375, *p* < 0.0001; *warbler:* mean ± s.e.= -1.757 ± 0.609, *cuckoo:* mean ± s.e.= 5.096 ± 0.821) than the cuckoo eggs in the phenotypic space.

**Figure 2.**
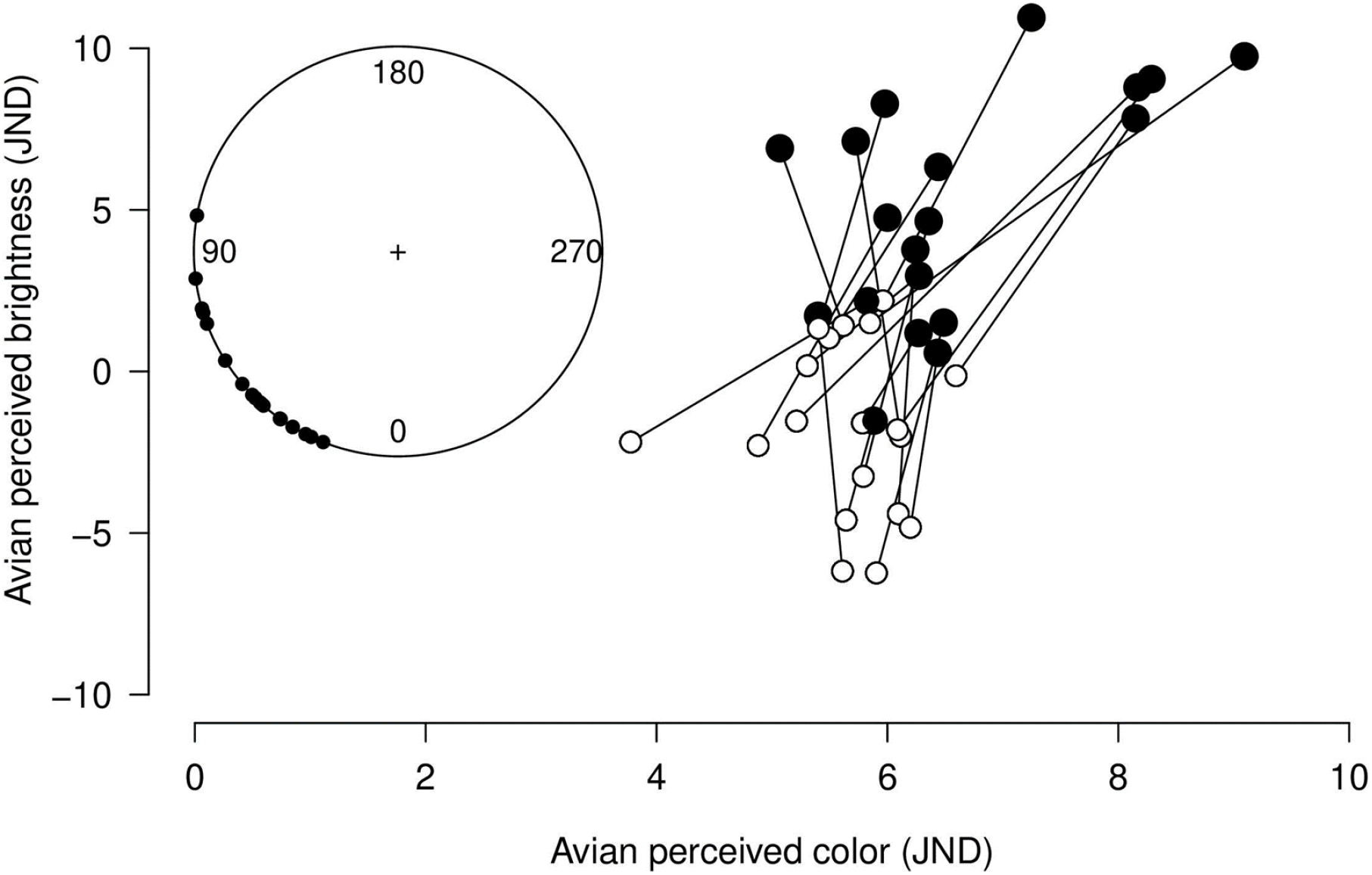
We estimated the Euclidean distances (solid lines) for the egg phenotypes of the great reed warblers (open dots) parasitized by the common cuckoo (closed dots) in the Czech host-parasite system. The egg phenotype of warblers and cuckoos is consistently different in avian perceived color (*x-axis*) and brightness (*y-axis*). Specifically, the warbler eggs are bluer and darker than the cuckoo eggs. The inset depicts the angles calculated for every host-parasite pair (close dots) plotted in a circular plane.

We found a significant nonuniform dispersion of the angles between host-parasite pairs toward a consistent direction on a 360º plane (*r* = 0.9347, *p* < 0.001; Fig. 2), suggesting a non-random direction of selection on the warblers by the cuckoos. These patterns were consistent even when we bootstrapped 10000 times for unique host-parasite combinations (mean ± s.e.= *r*: 0.928 ± 0.0003; *p*: 1.173e-7 ±1.621e-09; Fig. S2). As expected, we found that warblers were biased toward bluer and darker phenotypes; thus, the direction between the warbler and cuckoo populations is biased for the blue side of the avian eggshell color gradient.

## Discussion

The co-evolved eggshell mimicry between the eggs of the great reed warbler and its parasite, the common cuckoo, is a classic example of the coevolutionary arms races [19]. In well-established populations, a history of co-evolutionary arms races results in eggshell mimicry. Thus, in populations with long histories of host-parasite interaction, we would expect few systematic differences between the eggshell phenotypes of hosts and their parasites [7,32]. However, here we examined eggshell phenotypes within a phenotypic space that quantified variation in both coloration and brightness and found systematic differences between the eggshell phenotypes of hosts and their parasites. By doing so, we found non-overlapping eggshell phenotypes between hosts and parasites (see electronic supplementary material, figure S3), in congruence with prior observations that *Acrocephalus* warblers responded to both color and brightness differences [15]. These findings suggest that these co-evolutionary dynamics may be mediated through negative frequency-dependence selection [33], rather than via the classically accepted coevolutionary arms races [34,35].

Under co-evolutionary arms races, hosts adapt incrementally improved egg discrimination abilities, while the parasite adapts improved eggshell color mimicry [8,36]. As the host population purges the parasite eggs that are the poorest matches to their own, they select for eggshell mimicry within the parasite population. Thus, the eggshell mimicry between both populations improves over time [7]. After mimicry has evolved, the egg coloration of the host and parasite should overlap in a phenotypic space. Then, we expect no systematic differences in egg appearances between host-parasite pairs. By contrast, red queen dynamics are antagonistic interactions of negative frequency-dependent selection between hosts and parasites [37,38]. Such reciprocal selection causes fluctuations in the phenotypic frequencies of both parties [33,39]. Under this model of coevolution the host and parasite eggs would differ in appearance within the phenotypic space, since each generation will present new adaptations and counter-adaptations in eggshell appearance [32]. The host rejection selects for mimicry (i.e., good parasite matches as under arms races); but this in turn favors host eggshell phenotypes that remain differentiated from their parasite’s phenotype (e.g., both populations could evolve ever-bluer eggs) and against the most dissimilar parasite phenotypes. The parasites instead are rejected at a higher rate by hosts that present poor matches; therefore, the poor host matches are in advantage and would grow more common for a further generation [32]; ultimately, these dynamics lead to constant oscillations in the eggshell coloration over time for both parties [40] where parasites would closely track their hosts phenotype over evolutionary time [32].

Eggshell mimicry evolved by way of coevolutionary arms races will eventually reach an endpoint when the cognitive and physiological limits on the hosts (egg recognition) and parasites (egg pigment production) are met [41–43]. However, the integration of new phenotypes into a host-parasite system via migration or mutation might forestall the evolution of exquisite eggshell mimicry [44]. Due to these processes, it might be expected that mimicry would be imperfect even if the populations had long coevolutionary histories. Little is known on the recruitment of young cuckoos [but see, 45], although once recruited as adults, cuckoo females are known to stay within the same populations within a single season and across seasons [46]. Because host-parasite systems are dynamic, red queen dynamics might occur at any point in the coevolutionary process. These fluctuations of phenotypic frequencies over time may explain the diversification in egg appearance found in some systems [32,39,40,47].

Importantly, co-evolutionary arms races and red queen dynamics are neither mutually exclusive nor opposing dynamics [48]. Any moment of a host-parasite coevolution history can be governed by either these dynamics, or at an intermediate stage between these two processes [43,49,50]. Moreover, a wide range of population-level changes may push the host-parasite community towards a fluctuating dynamic where color-variants of parasite eggs can thrive even when they are dissimilar to the hosts eggs. Likewise, color-biased rejection behavior in these warblers would lead to differential survival based on egg phenotype [15], for both their parasite and (to a lesser extent) themselves. In fact, considering these factors it may be reasonable to assume that most established host populations experience some combination of both coevolutionary processes. Here we show that the eggs of great read warblers and common cuckoos from a specific population in Czech Republic have consistent differences in both avian perceived color and brightness; these differences indicate that eggshell perfect mimicry has not been met yet in this system. In combination with behavioral responses [15], these patterns suggest that eggshell cuckoo mimicry may be driven by a red queen dynamic, rather than via co-evolutionary arms races. Future research integrating multiple generations for the same host-parasite system would provide insights into the egg phenotype evolution between warblers and cuckoos.

## Supporting information

Electronic supplementary material

## Data Accessibility

All data and code will be publicly accessible via Dryad after pre-print submission.

## Authors’ Contribution

D.H. conceived the study; D.H., J.V., and P.W. planned the analyses; D.H., P.W. and J.V. generated and ran the analyses; J.V. and D.H. wrote the original draft. All authors approved publication.

## Acknowledgements

We thank Alex J. Di Giovanni for edits and comments on this paper. We also would like to thank the AOS, Blandy Experimental Farm, and the National Science Foundation and the College of Science at George Mason University for providing funds and space for developing this project.

