## Supplementary material for "Consistent differences in eggshell phenotypes select for bluer eggs in an avian host-parasite system": Electronic supplementary material

**Electronic Supplementary Material** Accompanies the manuscript:

**Supplementary Methods**

(a) *Phenotypic space*

We constructed a 2-dimensional phenotypic space where distances between coordinates correspond with both perceived variation in color and perceived variation in luminance of eggshell stimuli [1]. Here our goal is to compare eggshell phenotypes of hosts and parasites, but since the tetrahedral color space omits luminance information [2,3], such comparisons would be challenging within that space. However, we overcame this challenge by using a phenotypic space that considers both avian perceived color and brightness are integrated in a single plane [1,4]. Specifically, the x- and y-axis of the phenotypic space are represented as directional just noticeable differences (directional JNDs) of avian perceived color and avian perceived brightness, respectively (see main text for a complete description). Therefore, within this space we can compare eggs of the common cuckoo (*Cuculus canorus*), and its host the great reed warbler (*Acrocephalus arundinaceus*), relative to a common reference point. In this case, we chose the American robin (*Turdus migratorious*), which has eggs pigmented only by biliverdin. For convenience hereafter we will refer these species as ‘warbler’, ‘cuckoo’ and ‘robin’ respectively.

i. *Comparing positions within the phenotypic space*

To test our hypothesis, we needed to determine whether cuckoo eggshells were well matched to their hosts (i.e., had similar degrees of perceived color and brightness) or whether there were consistent differences between the parasites’ eggshell colors and their hosts’ eggshell colors (i.e., were bluer, browner, brighter, and/or darker). Here we needed to determine the angles between eggshell coordinates within this phenotypic space (main text, figure 1). The robin served as a common reference point to ensure that our angles were comparable within the phenotypic space. Although this could be the egg phenotype of any species’ egg, we selected the robin egg because its blue-green eggshell is mainly pigmented by biliverdin [5]. Since eggshell colors fall within a line in the avian visual color space, the chromatic differences to the robin’s eggshell color were highly related to the x-coordinate for the avian color space (Pearson’s r=0.906, IC_95_ = 0.825 to 0.950, t= 12.825, d. f.= 36, p < 0.001). Here, both warbler and cuckoo eggs from this Czech system should be browner than the robin egg. We collected fresh eggshell reflectance spectra of robin eggshells at the Blandy Experimental Research Farm in Boyce, Virginia, United States (39°3' N, 78°3' W). We measured the eggshell reflectance with a field portable spectrophotometer (Jaz Ocean Optics) and measured the spectra relative to a diffuse reflectance standard Ocean Insight (WS-1-SL), and to darkness at a coincident normal measurement angle. We took six measurements per egg (n=7), two per each of the three egg regions: blunt end, equator, and pointed end [6]. We then calculated the average reflectance spectra per- and between the robin eggs using the package in *pavo* in R (figure S1).

ii. *Calculating and quantifying angles*

First, we estimated Euclidean distances between the directional JNDs of each pair of cuckoo and warbler eggs, then between each cuckoo and warbler to the common reference point (vectors a, b, and c from Figure 2). We then estimated the angle formed between the coordinates of each host-parasite pair by applying the law of cosines,

$\cos C= \frac{a^{2}+b^{2}-c^{2}}{2ab}$	eq. 1

where *a*, *b* and *c* are respectively the Euclidean distances between the cuckoo and the warbler, the cuckoo and the robin, and the robin and the warbler (main text, figure 1). Here, C could be identical for warbler eggs falling above or below vector *b*; therefore, whenever the warbler egg phenotype has an avian perceived brightness greater than the slope intercept of the robin-cuckoo vector *b*. Conversely, we multiplied by -1 the angles were the warbler egg have a smaller value for the avian perceived brightness relative to the slope intercept of the vector *b.*

iii. *Phenotypic overlap between the egg of the great reed warbler and the common cuckoo*

To estimate the phenotypic overlap between warblers and cuckoos, we calculated the convex hull encompassing the coordinates for both the host and parasite within the phenotypic space [7]. The convex hull displays the smallest convex polygon that encloses all the points for each of the sets of data. Using these convex hulls, we counted the number of cuckoo eggs that fall into the warbler’s convex hull (i.e., within the intersection). Specifically, we found that 15.79% of the host (n=3) and 5.26% of the cuckoos (n=1) had a phenotype that fall into the cuckoo-host intersection (figure S3).

(b) *Comparing analyses: blue-green chroma and avian perceived differences*
i. *Color metrics*

To ensure our analyses were robust to different colorimetric scores, we repeated our analyses using standard colorimetric variables, specifically blue-green chroma and brightness [8]. We calculate the standard colorimetric variables, we estimated the blue-green chroma (BGC) as the proportion of total reflectance in the blue-green region of the spectrum,

$BGC=\frac{\sum_{450}^{550} R}{\sum_{300}^{700} R}$	eq. 2

where R represents reflectance [8]. We scaled from proportions to percentages the BGC values in order to make them comparable to the mean brightness, calculated as the total brightness relative to the number of wavelength intervals [8],

$\frac{\sum_{300}^{700} R}{n_{\lambda}}$	eq. 3

We then constructed a similar phenotypic space using blue-green chroma and mean brightness for each egg. The blue-green chroma is negatively correlated to JNDs of avian perceived color, while mean brightness was positively correlated to avian perceived brightness (table S1). Both correlations were significant and indicate that the phenotypic space can be created in terms of either standard colorimetric variables or perceptual differences. Finally, we calculated the angles between each warbler-cuckoo pair as outlined above. We then implemented circular statistics [9], test for uniformity using a Rayleigh’s test and determined that the angles between warbler-cuckoo pairs were uniformly distributed (*r* = 0.96, *p* < 0.001). We concluded that either standard colorimetric variables or perceptual model are reliable and comparable approaches to analyze phenotypic distances. Finally, constructing our phenotypic space using the x-coordinate within the avian color space (defining egg colors) and avian perceived luminance, produced nearly identical results (*data not shown*).

(c) *Test for pseudo replication*

Because female common cuckoos tend to parasitize nests that are located close to each other [10–12], we tested whether pseudo-replication could affect the uniformity and significance of the phenotypic analyses. To do so, we formed arbitrary host-parasite pairs between warblers and cuckoos (n=19). Then we repeated our approach (outlined above) to determine whether there were any consistent directional differences between cuckoos and their hosts (see ‘Methods’ in the main text). We repeated this step 10,000 times. We found that even after randomly mixing warblers and cuckoos, the values for the Rayleigh *r* statistics were uniform in the distribution of the host-parasite pairs within the phenotypic space (mean ± se: *r*=0.928 ± 0.0003, *p*=1.173e-07 ± 1.621e-09; figure S2).

**Table S1.** Pearson’s correlations between the color and brightness metrics of two different approaches: standard colorimetric variables and the receptor-noise limited model (just noticeable differences, JNDs). For each correlation we reported the Pearson’s correlation coefficient (r), the lower and upper 95% confidence limits (lcl and ucl), the *t* statistic (t), degrees of freedom (d. f.) and the significance (p).

| **Species** | **Correlation** | **r** | **lcl** | **ucl** | **t** | **d. f.** | **p** |
| --- | --- | --- | --- | --- | --- | --- | --- |
| Great reed warbler | *blue-green chroma ~ avian perceived color* | | | | | | |
|  |  | -0.937 | -0.976 | -0.841 | -11.066 | 17 | < 0.001 |
|  | *mean brightness ~ avian perceived brightness* | | | | | | |
|  |  | 0.968 | 0.916 | 0.988 | 15.867 | 17 | < 0.001 |
| Common cuckoo | *blue-green chroma ~ avian perceived color* | | | | | | |
|  |  | -0.796 | -0.918 | -0.536 | -5.432 | 17 | < 0.001 |
|  | *mean brightness ~ avian perceived brightness* | | | | | | |
|  |  | 0.984 | 0.957 | 0.994 | 22.62 | 17 | < 0.001 |

reflectance (%)

| wavelength (nm) | dat |
| --- | --- |
| 300 | 5.82343 |
| 301 | 6.0275 |
| 302 | 6.21002 |
| 303 | 6.43579 |
| 304 | 6.60277 |
| 305 | 6.73808 |
| 306 | 6.84744 |
| 307 | 6.93512 |
| 308 | 7.03352 |
| 309 | 7.15039 |
| 310 | 7.2445 |
| 311 | 7.31487 |
| 312 | 7.35774 |
| 313 | 7.40717 |
| 314 | 7.43249 |
| 315 | 7.43457 |
| 316 | 7.42707 |
| 317 | 7.40007 |
| 318 | 7.39433 |
| 319 | 7.35725 |
| 320 | 7.32447 |
| 321 | 7.28262 |
| 322 | 7.18574 |
| 323 | 7.08377 |
| 324 | 7.00787 |
| 325 | 6.90739 |
| 326 | 6.8594 |
| 327 | 6.85454 |
| 328 | 6.82847 |
| 329 | 6.76718 |
| 330 | 6.71432 |
| 331 | 6.58203 |
| 332 | 6.48797 |
| 333 | 6.43903 |
| 334 | 6.31044 |
| 335 | 6.25895 |
| 336 | 6.17605 |
| 337 | 6.05696 |
| 338 | 5.9745 |
| 339 | 5.84017 |
| 340 | 5.6745 |
| 341 | 5.56142 |
| 342 | 5.43713 |
| 343 | 5.35162 |
| 344 | 5.23999 |
| 345 | 5.10674 |
| 346 | 5.01433 |
| 347 | 4.90317 |
| 348 | 4.81413 |
| 349 | 4.73255 |
| 350 | 4.64625 |
| 351 | 4.57551 |
| 352 | 4.50506 |
| 353 | 4.39673 |
| 354 | 4.29101 |
| 355 | 4.18914 |
| 356 | 4.10683 |
| 357 | 4.04977 |
| 358 | 3.99804 |
| 359 | 3.93723 |
| 360 | 3.87303 |
| 361 | 3.81377 |
| 362 | 3.78811 |
| 363 | 3.7889 |
| 364 | 3.82179 |
| 365 | 3.86883 |
| 366 | 3.91408 |
| 367 | 3.94457 |
| 368 | 3.94772 |
| 369 | 3.95946 |
| 370 | 3.94226 |
| 371 | 3.95605 |
| 372 | 4.00523 |
| 373 | 4.02822 |
| 374 | 4.09455 |
| 375 | 4.1481 |
| 376 | 4.16459 |
| 377 | 4.21301 |
| 378 | 4.2553 |
| 379 | 4.32557 |
| 380 | 4.40875 |
| 381 | 4.49733 |
| 382 | 4.58649 |
| 383 | 4.64936 |
| 384 | 4.73955 |
| 385 | 4.83542 |
| 386 | 4.94076 |
| 387 | 5.0638 |
| 388 | 5.16619 |
| 389 | 5.28518 |
| 390 | 5.39122 |
| 391 | 5.48374 |
| 392 | 5.58965 |
| 393 | 5.70156 |
| 394 | 5.84226 |
| 395 | 5.99628 |
| 396 | 6.15967 |
| 397 | 6.31752 |
| 398 | 6.46942 |
| 399 | 6.63402 |
| 400 | 6.80611 |
| 401 | 6.99815 |
| 402 | 7.19662 |
| 403 | 7.377 |
| 404 | 7.551 |
| 405 | 7.72076 |
| 406 | 7.90085 |
| 407 | 8.11276 |
| 408 | 8.34669 |
| 409 | 8.55954 |
| 410 | 8.79793 |
| 411 | 9.0446 |
| 412 | 9.28928 |
| 413 | 9.58239 |
| 414 | 9.87403 |
| 415 | 10.1439 |
| 416 | 10.4243 |
| 417 | 10.7097 |
| 418 | 10.9912 |
| 419 | 11.285 |
| 420 | 11.5732 |
| 421 | 11.8763 |
| 422 | 12.1941 |
| 423 | 12.5315 |
| 424 | 12.8861 |
| 425 | 13.2314 |
| 426 | 13.5947 |
| 427 | 13.9871 |
| 428 | 14.3936 |
| 429 | 14.824 |
| 430 | 15.2352 |
| 431 | 15.6407 |
| 432 | 16.0538 |
| 433 | 16.4526 |
| 434 | 16.8418 |
| 435 | 17.242 |
| 436 | 17.6472 |
| 437 | 18.0681 |
| 438 | 18.4764 |
| 439 | 18.8691 |
| 440 | 19.255 |
| 441 | 19.6327 |
| 442 | 20.0138 |
| 443 | 20.4179 |
| 444 | 20.8352 |
| 445 | 21.2258 |
| 446 | 21.6077 |
| 447 | 21.9658 |
| 448 | 22.325 |
| 449 | 22.7088 |
| 450 | 23.0903 |
| 451 | 23.4628 |
| 452 | 23.8241 |
| 453 | 24.1699 |
| 454 | 24.5134 |
| 455 | 24.8646 |
| 456 | 25.2001 |
| 457 | 25.5056 |
| 458 | 25.8015 |
| 459 | 26.0916 |
| 460 | 26.3647 |
| 461 | 26.6446 |
| 462 | 26.9338 |
| 463 | 27.2356 |
| 464 | 27.5281 |
| 465 | 27.8001 |
| 466 | 28.0816 |
| 467 | 28.3482 |
| 468 | 28.6071 |
| 469 | 28.871 |
| 470 | 29.1184 |
| 471 | 29.3418 |
| 472 | 29.5691 |
| 473 | 29.795 |
| 474 | 30.0166 |
| 475 | 30.2231 |
| 476 | 30.4246 |
| 477 | 30.6049 |
| 478 | 30.7669 |
| 479 | 30.9538 |
| 480 | 31.1138 |
| 481 | 31.2632 |
| 482 | 31.641 |
| 483 | 32.0918 |
| 484 | 32.36 |
| 485 | 32.5574 |
| 486 | 32.6141 |
| 487 | 32.5699 |
| 488 | 32.5829 |
| 489 | 32.6533 |
| 490 | 32.7297 |
| 491 | 32.8034 |
| 492 | 32.8969 |
| 493 | 32.9916 |
| 494 | 33.0853 |
| 495 | 33.1525 |
| 496 | 33.1906 |
| 497 | 33.2278 |
| 498 | 33.2785 |
| 499 | 33.3338 |

**Figure S1.** Reflectance spectrum of the American Robin egg.

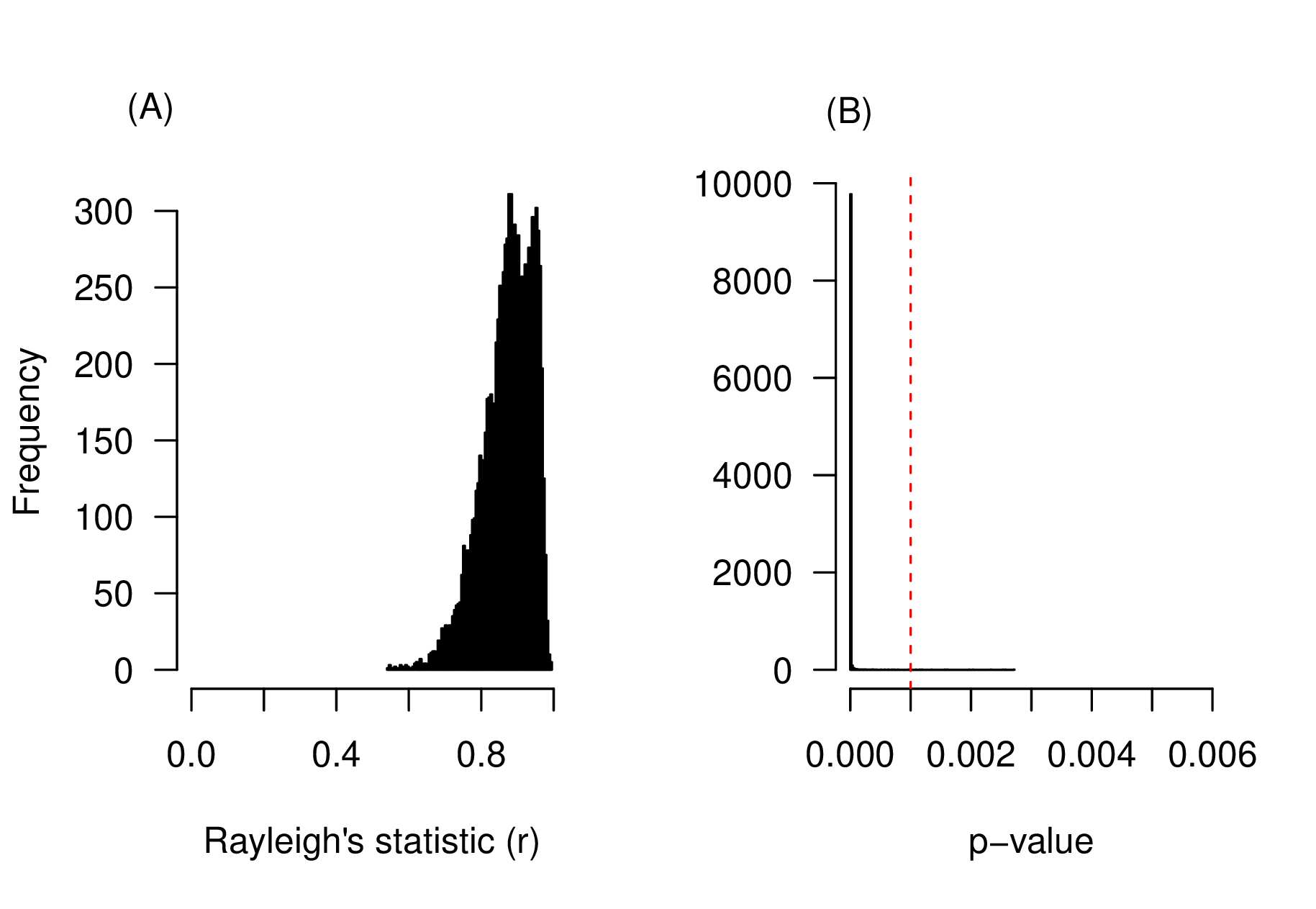

**Figure S2.** The frequencies of the Rayleigh’s test statistic r (A) and the p-value (B) after resampling 10000 times the warbler-cuckoo system of Czech Republic. The dashed red line denotes a p-value significance equal to 0.001.

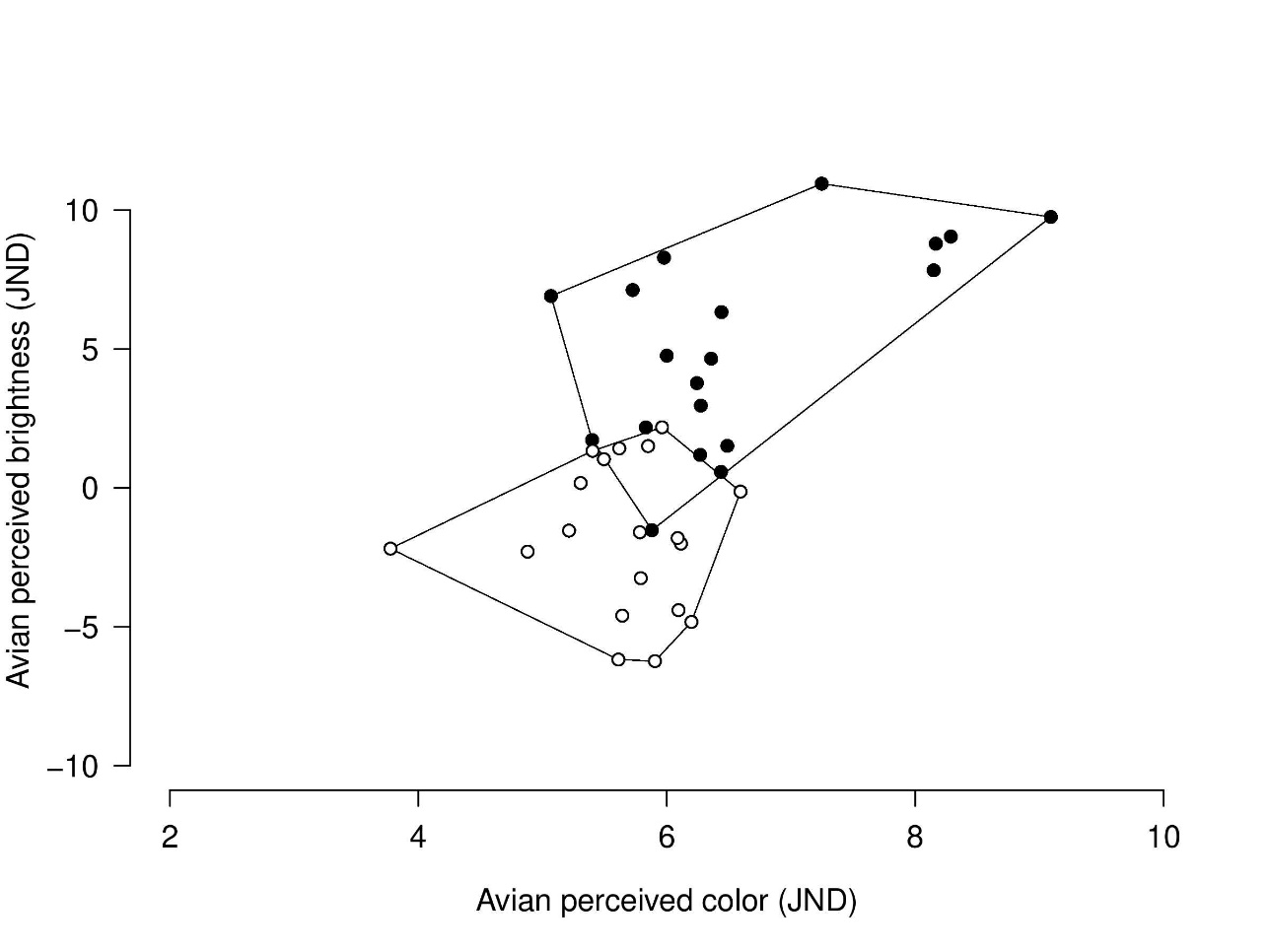

**Figure S3.** We display the convex hull polygons within the phenotypic space for the warbler (open dots) and the cuckoo (close dots) eggshell phenotypes.
